# A role for LGI1 in regulating pain sensitivity

**DOI:** 10.1101/2023.09.13.557645

**Authors:** Adham Farah, Ivan Paul, Hoi Cheng, Yuhe Su, Piotr Poplawski, Mandy Tseng, John Dawes

## Abstract

Chronic pain represents a major unmet clinical need. Neuropathic pain, that is pain arising due to damage or disease of the somatosensory nervous system, represents a sizeable proportion of chronic pain cases, affecting around 8% of the general population. Neuronal hyperexcitability is a key driver of neuropathic pain. Leucine glioma inactivated 1 (LGI1), is a secreted protein known to regulate excitability within the nervous system and is the target of autoantibodies from neuropathic pain patients. Therapies that block or reduce antibody levels are highly effective at relieving pain in these patients, suggesting that LGI1 has an important role in clinical pain. Here we have studied the role of LGI1 in regulating pain using mouse models to specifically ablate LGI1 in neuron populations. LGI1 has been well studied at the level of the brain. Here we show that LGI1 is highly expressed in dorsal root ganglion (DRG) neurons and in dorsal horn neurons of the spinal cord. Using transgenic mice, we ablated LGI1, either specifically in nociceptors (LGI1^fl/Nav1.8(+/-)^), or in all DRG and spinal neurons (LGI1^fl/Hoxb8(+/-)^). On acute pain assays, mild phenotypes were observed when compared to littermate controls with limited changes in DRG neuron excitability. No differences were seen in the first phase of the formalin test, however LGI1^fl/Hoxb8(+/-)^ mice displayed a significant increase in nocifensive behaviours in the second phase compared to littermate controls. Using the spared nerve injury model, we assessed the impact of LGI1 ablation on neuropathic pain-like behaviours. LGI1^fl/Nav1.8(+/-)^ mice showed no differences in nerve injury induced mechanical hypersensitivity, brush-evoked allodynia or spontaneous pain behaviour compared to controls. However, LGI1^fl/Hoxb8(+/-)^ mice showed a significant exacerbation of mechanical hypersensitivity and allodynia. These data show that LGI1 has a role in regulating pain sensitivity particularly in the context of nerve injury. We suggest this effect is likely mediated at the spinal level since no differences were observed following specific ablation of LGI1 in nociceptors. Neurons in dorsal horn of the spinal cord are important in gating nerve injury induced mechanical pain and our findings suggest that LGI1 plays an important role in this process.

## INTRODUCTION

Neuropathic pain, a long-lasting pain condition typically caused by a lesion or disease of the somatosensory nervous system, is estimated to have a prevalence in the general population of around 8% (Cavalli et al., 2019). Although analgesics are available, neuropathic pain is particularly resistant to treatment and current therapies cause adverse side effects, thus negatively affecting the overall quality of life in patients and leading to substantial economic and societal burdens (Fornasari, 2017; Gaskin & Richard, 2012). Excessive neuronal excitability is the basis of many persistent pain conditions such as neuropathic pain. There is now ample experimental and clinical evidence to support that both peripheral and central changes contribute to neuronal hyperexcitability in neuropathic pain. For instance, blocking the increased excitability of primary sensory neurons can alleviate evoked and ongoing pain in patients (Haroutounian et al., 2014). Spontaneous activity in DRG neurons following nerve injury, drives central changes leading to an enhanced response of dorsal horn neurons (Woolf, 2011). These activity depending changes are accompanied by a loss of inhibitory tone in the spinal cord further contributing to spinal hyperexcitability (Todd, 2015). Therefore a better understanding of the molecular pathways that influence neuron excitability in pain pathways will greatly assist in the development of new better targeted pain therapies.

Leucine glioma inactivated 1 (LGI1) is a secreted molecule important for protein interactions in the nervous system. It has a known role in driving hyperexcitability within the nervous system in the context of disease. For example, loss of function mutations are linked to epilepsy in humans (Kalachikov et al., 2002). In line with this, LGI1 has been shown to regulate synaptic transmission through its interaction and modulation of ion channels. For example, genetic ablation of LGI1 in mice leads to a loss of pre-synaptic Kv1 channels and a loss of AMPA receptors postsynaptically (Fukata et al., 2006; Schulte et al., 2006). It can also regulate the intrinsic excitability of neurons in a Kv1 dependent manner (Seagar et al., 2017). LGI1 is the target of autoantibodies in neuropathic pain patients and pain can be relieved in these patients with therapies that either block or reduce the level of LGI1 antibodies (Gadoth et al., 2017; Ramanathan et al., 2021). These observations suggest that disruption of LGI1 is sufficient to cause neuropathic pain in patients and that LGI1 plays a role in regulating pain sensitivity.

Here, we aim to better understand the role of LGI1 in regulating neuron excitability and pain sensitivity both in the acute and pathological setting, using conditional genetic ablation in mice.

## RESULTS

### LGI1 expression in DRG neurons and spinal cord

Using in situ hybridization (ISH) and immunohistochemistry (IHC) done on mouse primary sensory neurons from lumbar DRG, we found LGI1 signal in all DRG neuron subpopulations tested, including large myelinated, NF200-positive cells, CGRP-positive neurons and those that bind IB4 (Fig.1A, Table S1). In line with these findings and since LGI1 is a secreted protein, we also assessed the expression of known LGI1 interactors (ADAM11, 22, 23) in DRG neurons and found a similar expression profile to that of LGI1 with the highest level of ADAMs expression was in cells that express NF200 (Fig. S1A, B, Table S1).

**Fig. 1:**
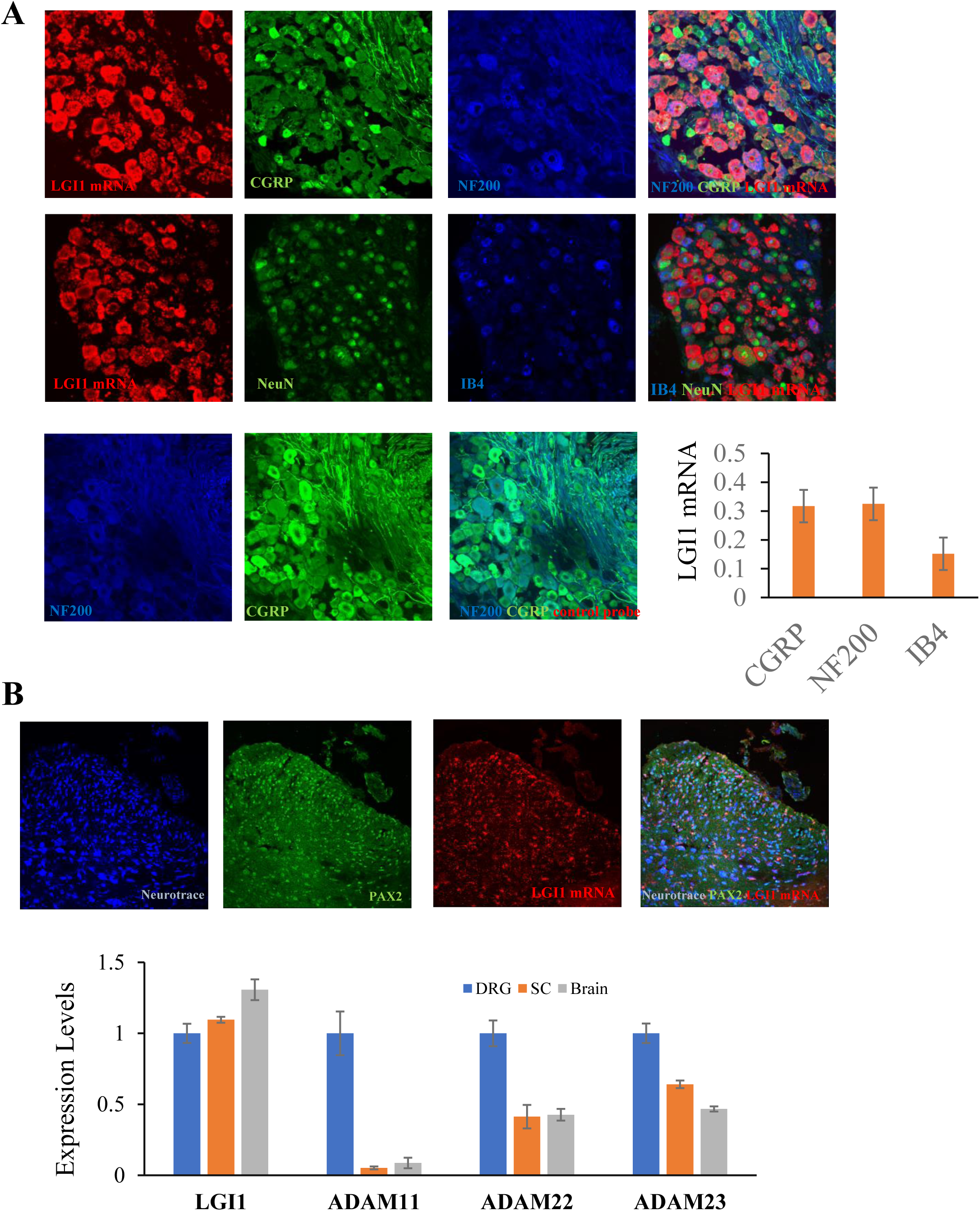
LGI1 expression in DRG and spinal cord. **(A)** LGI1 is expressed in DRG neurons (Lumbar 4). Representative images of ISH for LGI1 mRNA in DRG sections stained for molecular markers of DRG neuron subtypes: IB4, CRGP (nociceptors), and NF200 (large, myelinated touch fibres). Quantification of signal intensity for LGI1 probe. Data shown as mean±SEM (n=3-4 mice) **(B)** LGI1 expression in the mouse spinal cord. Representative images of ISH for LGI1 in spinal cord showing also neurotrace-positive neurons co-localizing with LGI1 & PAX2 in the superficial laminae and in the deeper laminae of the spinal cord. **(C)** qPCR data analysis also confirms the expression of LGI1 in DRG, spinal cord and brain with no significant differences for LGI1 expression across all three tissues, but ADAM11,22,23 showed lower expression levels in the spinal cord and brain relative to the DRG levels. Data shown as mean±SEM (n=3).

As detailed anatomical analysis of LGI1 expression in the mouse spinal cord is lacking, we conducted ISH which revealed that neurotrace-positive neurons co-localized with LGI1 in the superficial laminae and in the deeper laminae of the spinal cord. We also confirmed LGI1 expression within PAX2-positive inhibitory interneurons (Fig.1B). ISH images also indicate that ADAM 11, 22, and 23 co-localized with PAX2- and neuotrace-positive neurons (Fig. S1C).

We also measured the expression of LGI1 and ADAMs in the DRG, spinal cord, and brain using qPCR, and saw no significant differences for LGI1 expression across all three tissues, but the ADAMs showed lower expression levels in the spinal cord and brain relative to the DRG levels (Fig. 1C, Table S2).

We have also examined the expression levels of the proteins LGI2, 3, and 4 as these also play roles in the central and peripheral nervous system (Kegel et al., 2013). Our qPCR results indicate that LGI2 and 3 had a similar expression pattern and profile to that of LGI1 in both the DRG and spinal cord, and lesser expression levels were detected in the brain in comparison to LGI1 expression levels. On other hand, negligible expression levels were detected for the LGI4 protein in all three types of tissues relative to LGI1 expression levels (Fig. S3A, Table S3).

In summary, LGI1 is highly expressed both at the level of the DRG and spinal cord in mice, and its binding partners, ADAMs, are also widely expressed in the DRG and to a lesser extent in the spinal cord.

### LGI1 Regulates Pain-Related Hypersensitivity in Mice

First, we performed qPCR to confirm that LGI1 was knocked down at the level of the DRG in LGI1^fl/Nav1.8(+/-)^ vs control mice (LGI1^fl/Nav1.8(-/-)^-WT). The expression level of LGI1 was decreased by 70% in the LGI1 knockout mice (cKO) from the Nav 1.8 Cre mouse line (p<0.001, n=3, Fig. S2A). A similar validation for the HoxB Cre line was performed in which LGI1 expression was significantly reduced at all DRG and spinal levels (p<0.001, n=3, Fig. S2B,C, Table S4). We also examined the expression levels of the ADAMs and the proteins LGI2, 3, and 4 in the mouse line. Our qPCR data indicate that the expression levels of the ADAMs proteins were mostly similar between the cKO and WT mice in the DRG and spinal cord, apart from ADAM11 protein which showed a slight significant decrease in expression levels on the cervical level of the DRG and spinal cord of the cKO mice compared to the WT (p<0.05, n=3). The expression levels of LGI2, 3 and 4 were all similar between the two genotypes in the DRG and spinal cord (Fig. S3B & C, Table. S4).

Using our cKO lines, we then find that specific ablation of LGI1 in nociceptors results in a significant heat hypersensitivity vs WT littermate control (LGI1^fl/Nav1.8(-/-)^ 10.4±0.6s vs LGI1^fl/Nav1.8(+/-)^ 8.3±0.4s, n=14-21, Fig. 2A, Table S5) but not in the HoxB Cre line (p>0.05, n=19-22, Fig. 2B, Table S6). No significant differences were seen on the Hotplate (53°C) in both Cre lines (p>0.05, Fig. 2A, B, Table S5, 6). LGI1^fl/Nav1.8(+/-)^ mice did not show any difference to control in response to mechanical stimuli (Von Frey, pin prick, p>0.05, Fig. 2A, Table S5). On the other hand, LGI1^fl/Hoxb^ ^(+/-)^ mice were hypersensitive to Von Frey hair application, demonstrating a significantly lower withdrawal threshold compared to WT littermates (LGI1^fl/Hoxb^ ^(-/-)^ 0.61±0.02, n=25 vs LGI1^fl/Hoxb^ ^(+/-)^ 0.53±0.02; n=31; p<0.05; Fig. 2B, Table S6), but there were no differences between genotypes for the noxious pinprick application (p>0.05, Fig. 2B, Table S6).

**Fig. 2:**
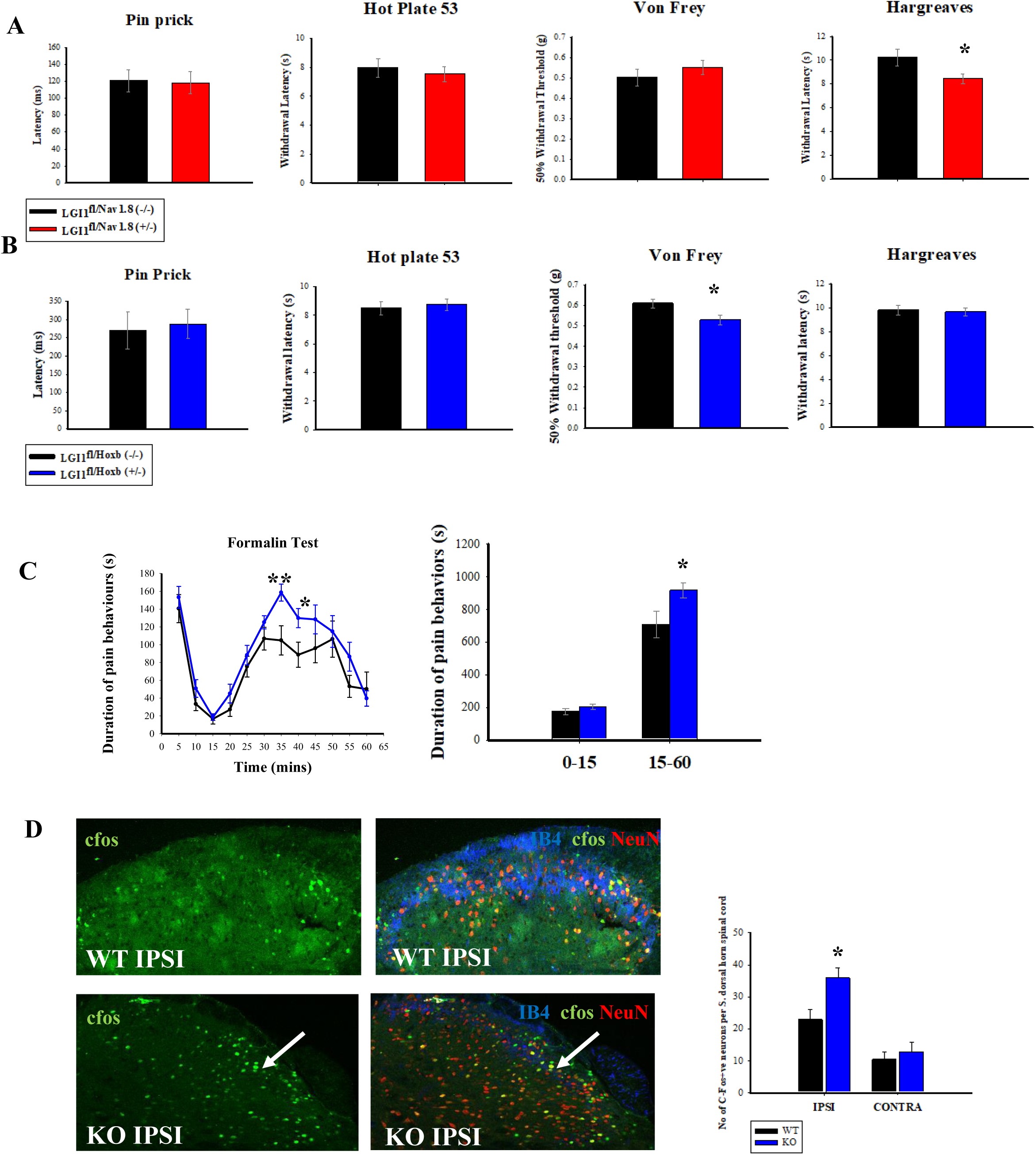
Sensitivity to mechanical (Von Frey, pin prick), thermal (Hot plate, Hargreaves) and chemical (formalin) stimuli was assessed in Nav1.8 Cre line & HoxB Cre line. **(A), (B)** Results show that nociceptor specific KO of LGI1 from LGI1^fl/Nav1.8(+/-)^ (n=21, *p<0.05, t test, mean±SEM) leads to heat hypersensitivity. Genetic ablation of LGI1 from LGI1^fl/HoxB(+/-)^ leads to mechanical hypersensitivity (n=31, *p<0.05, t test, mean±SEM) and pain hypersensitivity **(C)** during the second phase (15-60 min) of the formalin test when compared to littermate controls (n=14, *p<0.05, two-way repeated measures ANOVA posthoc Tukey or t test, mean±SEM). **(D)** Assessment of c-fos, a neural activity marker, in the superficial dorsal horn spinal cord did reveal a significant difference between genotypes (n=3, *p<0.05, One way ANOVA, mean±SEM).

We have also assessed sensitivity to chemical algogens specifically formalin application. In the HoxB Cre line, loss of LGI1 resulted in enhanced nocifensive responses during the second phase (15-60 min) of the formalin test compared to the control mice (LGI1^fl/Hoxb^ ^(-/-)^ 708.5±79.9s, n=10 vs LGI1^fl/Hoxb^ ^(+/-)^ 916.5±45.7s; n=14; p<0.05; Fig. 2C). Assessment of c-fos in the superficial layers of the spinal cord did reveal a significant difference between genotypes, in which the LGI1^fl/Hoxb^ ^(+/-)^ mice had a higher number of c-fos positive neurons (35.8±3.2; n=3) compared to the LGI1^fl/Hoxb^ ^(-/-)^ control mice (22.75±3.4; n=3; p<0.05, Fig. 2D), suggesting that the increased behavioral response in the second phase may be driven by a spinal mechanism.

### The effects of LGI1 on DRG soma sensitivity and excitability

We next used calcium imaging and cultured DRG neurons from control mice as well as cKOs, to assess the effect of disrupting LGI1 on DRG neuron sensitivity *in vitro*. Detection of any spontaneous activity was investigated as well as responses to chemical algogens such as capsaicin (1μM) and ATP (10μM) to activate specific nociceptor populations. No significant difference was observed between genotypes for the Nav 1.8 Cre line for spontaneous activity (n=6 mice, ∼1900 cells per genotype, p>0.05; Fig. S4A, Table S7). There was also no difference in the percentage of cells responding to either capsaicin or ATP application (p>0.05). Similar results were obtained for the HoxB Cre line, as there was no detected difference for the percentage of cells responding to ATP or capsaicin treatments as well for any spontaneous activity (n=3 mice, ∼500 cells per genotype, p>0.05; Fig. S4B, Table S7).

Using cultured primary sensory neurons from WT and LGI1 cKO mice, excitability parameters such as input resistance, rheobase and repetitive firing were assessed. No difference between the LGI1 cKO in both Cre lines and WT mice was observed for input resistance and rheobase (Small cells (<25 μm) LGI1^fl/Nav1.8(-/-)^ n=39, LGI1^fl/Nav1.8(+/-)^ n=35/ LGI1^fl/Hoxb^ ^(-/-)^ n=47, LGI1^fl/Hoxb^ ^(+/-)^ n=46; Medium cells (25–35 μm) LGI1^fl/Nav1.8(-/-)^, n=17 LGI1^fl/Nav1.8(+/-)^, n=20/ LGI1^fl/Hoxb^ ^(-/-)^ n=23, LGI1^fl/Hoxb^ ^(+/-)^ n=23; Fig.3A,B, Table S8). However, quantification across a range of current steps showed that in the HoxB Cre line small but not medium cKO diameter neurons display increased firing frequency in comparison to WT neurons (LGI1^fl/Hoxb^ ^(-/-)^: 1.15±0.17 vs LGI1^fl/Hoxb^ ^(+/-)^: 2.04±0.59; p<0.01; Fig 3B).

**Fig. 3:**
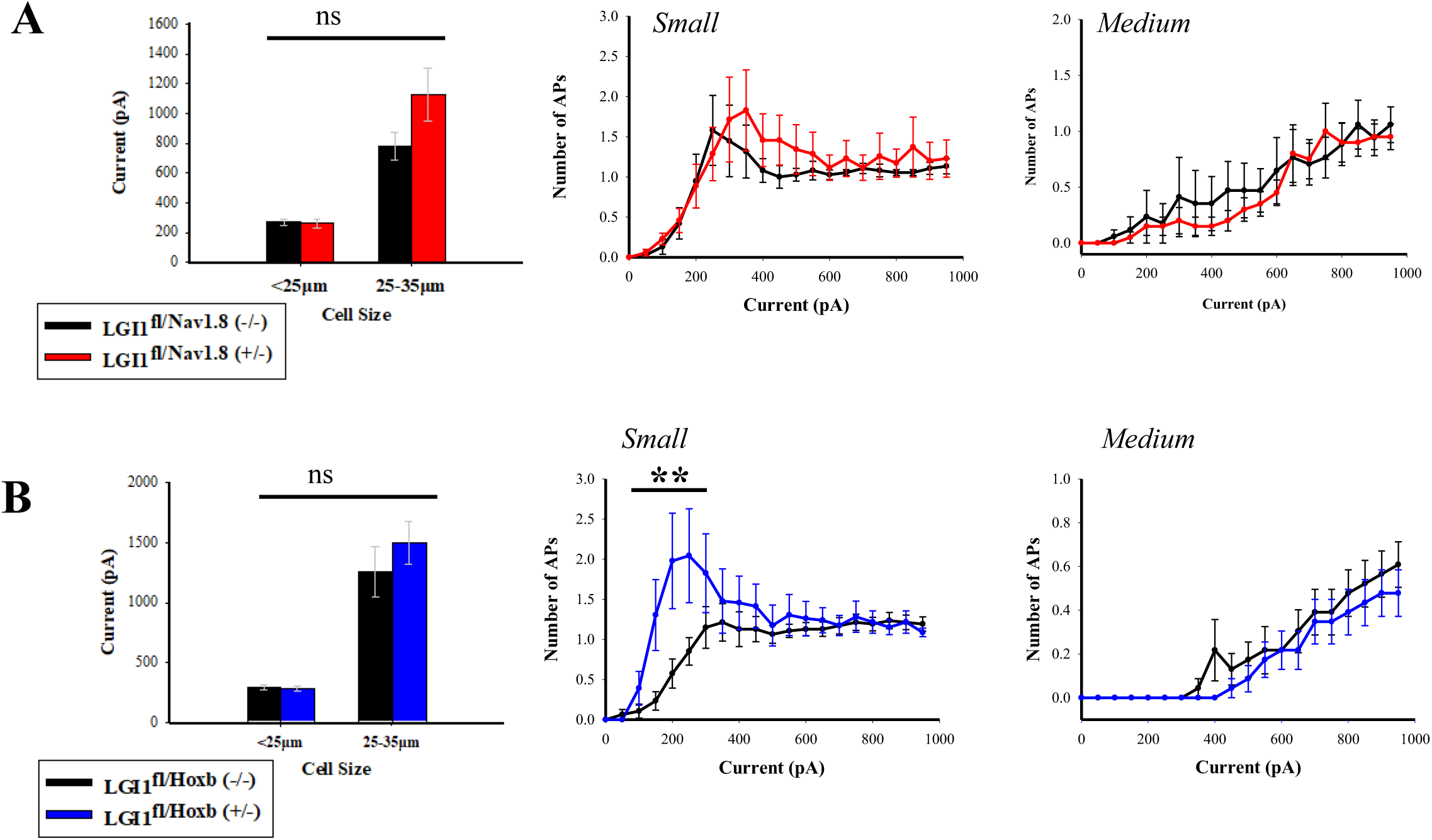
Using DRG neuron from LGI1^fl/Nav1.8(-/-)^ or LGI1^fl/HoxB(-/-)^ and LGI1^fl/Nav1.8(+/-)^ or LGI1^fl/HoxB+/-)^ mice, excitability parameters, rheobase and repetitive firing, were assessed in cultured primary sensory neurons. **(A)** No difference between the two genotypes for the Nav1.8 Cre line was observed for these excitability parameters (Small cells (<25 μm) LGI1^fl/Nav1.8(-/-)^ n=39 LGI1^fl/Nav1.8(+/-)^ n=35; Medium cells (25–35 μm) LGI1^fl/Nav1.8(-/-)^ n=17 LGI1^fl/Nav1.8(+/-)^ n=20). **(B)** Small (LGI1^fl/HoxB(-/-)^ n=47 LGI1^fl/HoxB+/-)^ n=46) and medium (LGI1^fl/HoxB-/-)^ n=23 LGI1^fl/HoxB(+/-)^ n=23) diameter DRG neurons cultured from HoxB Cre line showed no significant difference for rheobase when compared to neurons from control mice. However, quantification across a range of current steps showed that small LGI1^fl/HoxB(+/-)^ but not medium LGI1^fl/HoxB(+/-)^ diameter neurons display increased firing frequency in comparison to LGI1^fl/HoxB(-/-)^ neurons. **p<0.01 two-way repeated measures ANOVA posthoc Tukey, mean±SEM.

### Genetic ablation of LGI1 exacerbates nerve injury induced pain

To assess the effects of the genetic removal of LGI1 in the context of nerve injury, spared nerve injury (tibial) model (tSNI) was chosen as a model of neuropathic pain. All mice received surgery and the mechanical withdrawal thresholds of mice and their thermal sensitivity on the ipsi side were measured. We first tested the LGI1^fl/Nav1.8(-/-)^ and LGI1^fl/Nav1.8(+/-)^ mice, and all mice became hypersensitive to mechanical stimuli and cold stimuli by day 3 and 6, respectively (p<0.001 vs baseline, n= 7 for LGI1^fl/Nav1.8(-/-)^ and n= 8 for LGI1^fl/Nav1.8(+/-)^), but no significant differences between genotypes were found (p>0.05, Fig. S5A-C). For the HoxB Cre line, both LGI1^fl/Hoxb^ ^(-/-)^ and LGI1^fl/Hoxb^ ^(+/-)^ mice developed mechanical and thermal hypersensitivity by day 3 and 6, respectively (p<0.001 vs baseline, n=15 for LGI1^fl/Hoxb^ ^(-/-)^ and n=17 for LGI1^fl/Hoxb (+/-)^, Fig. 4A -D). However, the LGI1^fl/Hoxb^ ^(+/-)^ mice had a significantly higher hypersensitivity to Von Frey after tSNI from day 3 (Day3: LGI1^fl/Hoxb^ ^(-/-)^: 0.35±0.07 vs LGI1^fl/Hoxb^ ^(+/-)^: 0.16±0.04; p<0.05) and to brush stimulation on days 14 and 28 compared to LGI1^fl/Hoxb^ ^(+/-)^ mice (Day 14: LGI1^fl/Hoxb^ ^(-/-)^: 1.07±0.11 vs LGI1^fl/Hoxb^ ^(+/-)^: 1.37±0.08; p<0.05). Since there is also a significant difference in the baseline measurements for the Von Frey test, the percentage of change from baseline was calculated which also indicated that LGI1 cKO mice were more hypersensitive than WT from day 3 after tSNI (Day14: LGI1^fl/Hoxb^ ^(-/-)^: 49.28±8.1 vs LGI1^fl/Hoxb^ ^(+/-)^: 77.80±2.62; p<0.01, Fig.4C). As for thermal sensitivity assessed by the cold preference test, no significant differences between genotypes were found (p>0.05, Fig.4D, S5C).

**Fig. 4:**
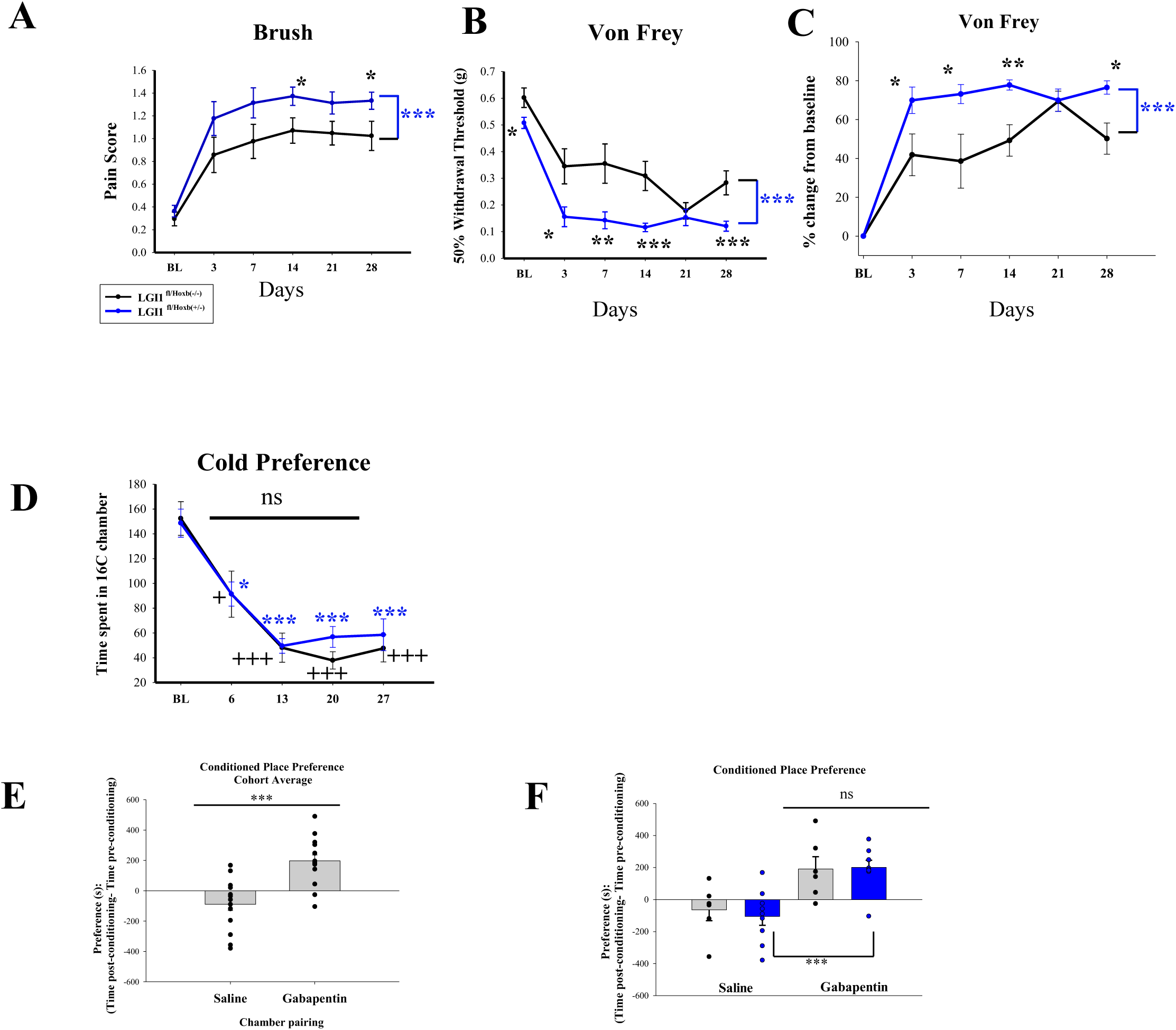
The effects of the genetic removal of LGI1 in the context of tSNI were assessed in the HoxB Cre line. **(A-D),** both LGI1^fl/Hoxb^ ^(-/-)^ (n=15) and LGI1^fl/Hoxb^ ^(+/-)^ (n=17) mice developed mechanical and thermal hypersensitivity after tSNI (for cold preference test: * vs baseline (BL), + vs BL, * and + p<0.05, *** and +++ p<0.001, two-way repeated measures ANOVA posthoc Tukey, mean±SEM). However, the LGI1^fl/Hoxb^ ^(+/-)^ mice had a significantly higher hypersensitivity to brush stimulation **(A)** and to Von Frey after tSNI on ipsi sides compared to LGI1^fl/Hoxb^ ^(-/-)^ mice, with percent change from baseline also calculated **(B & C)** (*<0.05, **p<0.01, ***p<0.001, two-way repeated measures ANOVA posthoc Tukey, mean±SEM). **(E-F)** ongoing neuropathic pain was examined in mice following tSNI using the conditioned place preference test, and gabapentin produced significant preference for the chamber paired with the drug **(E)**, however, no significant differences in the CPP difference scores between genotypes **(F)** were observed (LGI1^fl/HoxB(-/-)^ (n=6) and LGI1^fl/HoxB(+/-)^ (n=9), ***p<0.001, t test or One Way ANOVA, mean±SEM).

To assess ongoing pain, we used the conditioned place preference (CPP) paradigm, where the analgesic gabapentin or vehicle control were paired to different chambers (Griggs et al., 2015). We first tested the Nav1.8 Cre line mice, which showed an increased trend in preference for the gabapentin paired chamber, and the difference in CCP difference scores between the genotypes were nonsignificant (p>0.05, n= 7 for LGI1^fl/Nav1.8(-/-)^ and n= 8 for LGI1^fl/Nav1.8(+/-)^, Fig. S5D). For the HoxB Cre line, this dose of gabapentin produced significant preference for the chamber paired with the drug (gabapentin preference time: 197.3±38.92 vs saline preference time: -88.8±41.60, p<0.001), revealing gabapentin’s efficacy in relieving ongoing neuropathic pain, however, no significant differences in the CPP difference scores between genotypes were observed (p>0.05, n=6 for LGI1^fl/Hoxb^ ^(-/-)^ and n=9 for LGI1^fl/Hoxb^ ^(+/-)^, Fig. 4E,F).

## DISCUSSION

LGI1 is known to regulate excitability within the nervous system and is the target of autoantibodies from neuropathic pain patients. Here, using genetic ablation models, we find that LGI1 has a role in regulating pain particularly in the pathological setting.

LGI1 has been well studied at the level of the brain where it is genetically linked to epilepsy in patients (Kalachikov et al., 2002). However, studies are lacking on its role in the PNS and at the level of the spinal cord. We find that LGI is highly expressed by DRG and spinal cord dorsal horn neurons in mice. This is in line with recent single cell sequencing data (Häring et al., 2018; Zeisel et al., 2018). LGI1 is known to interact with ADAM proteins (11, 22 and 23) (Sagane et al., 2008), a key step in exerting its functional influence on the nervous system. Importantly we find that ADAM proteins are also well expressed both in the DRG and spinal cord. Therefore, we decided to remove LGI1 from these neuron populations and assess the impact on pain sensitivity. Due to the high expression of LGI1 in the DRG and spinal cord, we used the HoxB8 Cre to remove LGI1 in all lumbar DRG and spinal neurons and compare results to specific ablation from nociceptors using the Nav1.8 Cre line (Stirling et al., 2005). We found subtle effects on acute pain behaviour. For example, mild heat hypersensitivity in the LGI1^fl/Nav1.8(+/-)^ line and mechanical hypersensitivity in the LGI1^fl/Hoxb(+/-)^ line when compared to littermate controls. We next looked at the impact of LGI1 ablation on DRG neuron activity. Using calcium imaging and whole cell patch clamp electrophysiology, we saw no differences in the excitability of DRG neurons from LGI1^fl/Nav1.8(+/-)^. When LGI1 was removed from all DRG neurons, again no differences in activity were seen using calcium imaging and action potential thresholds were normal. However, a significant increase in repetitive firing was found in small DRG neurons. Genetic ablation of LGI1 increases the excitability of hippocampal neurons through its modulation of Kv1 channels (Seagar et al., 2017) and Kv1 channels have been shown to regulate repetitive firing in DRG neurons (Zhao et al., 2013). Therefore, the repetitive firing seen here is likely mediated through similar mechanisms. However, despite its high expression in DRG neurons, the effects on acute pain and excitability were subtle. This could be due to compensatory mechanisms, for example we have previously shown that CASPR2, also a target of autoantibodies in pain patients, is a strong regulator of Kv1 channel function in DRG neurons (Dawes et al., 2018). Alternatively, LGI1 may not play an important role in regulating excitability at this level. In line with this, we found that in the formalin model genetic ablation in both DRG and spinal cord did not affect the first phase of pain behaviour attributed to primary sensory neuron activity. However, a significant increase in pain behaviour was observed in the second phase strongly suggestive of a spinal role of LGI1 in regulating pain.

Since antibodies against LGI1 are associated with neuropathic pain in patients, we next wanted to assess the impact of LGI1 ablation on pain behaviours following nerve injury. Compared to littermate controls we found that ablation of LGI1 from nociceptors had no impact on mechanical or cold hypersensitivity or spontaneous pain following nerve injury. However, LGI1^fl/Hoxb(+/-)^ mice showed robust exacerbation of nerve injury induced mechanical pain hypersensitivity and brush evoked allodynia. Since nociceptor KO of LGI1 did not affect pain behaviours in the tSNI, our findings point to a spinal role of LGI1 in this context. Interestingly, in comparison to CASPR2-Ab patients, the distribution of pain in LGI1-Ab patients is more truncal suggestive of spinal involvement (Ramanathan et al., 2021). The exact mechanisms by which loss of spinal LGI1 enhances neuropathic pain remains to be investigated. However, it is likely to involve regulation of spinal neuron excitability through its modulation of Kv1 channels and AMPA receptors (Fukata et al., 2006; Schulte et al., 2006). In the context of epilepsy, LGI1 modulation of Kv1 and AMPA receptors have been attributed to the hyperexcitability and seizure development seen following global LGI1 ablation (Chabrol et al., 2010). Interestingly, a loss of AMPA receptor expression following LGI1 KO is thought to occur on GABAergic neurons resulting in a loss of inhibitory control (Fukata et al., 2017, 2021), reminiscent of the loss of inhibitory tone seen in the spinal cord following nerve injury (Todd, 2015). Nerve injury induced mechanical pain hypersensitivity, in particular touch evoked allodynia is known to be mediated at the level of the spinal cord (Boyle et al., 2019; Cheng et al., 2017). Recent studies have shed light on the neuronal populations involved and LGI1 may represent an important molecule underlying excitability change within these circuits. While future studies are needed to provide additional mechanistic insight, our findings point to a novel and important role for spinal LGI1 in regulating pain sensitivity particularly in the context of nerve injury.

## MATERIAL AND METHODS

### Mouse lines and animal care

All procedures were carried out in accordance with UK home office regulations and in line with the Animals Scientific Procedures Act 1986 at a licensed facility within the University of Oxford, following institutional review board approval. Animals were group housed in IVC cages in temperature and humidity-controlled rooms where food and water were available ad libitum, with a 12-hour light dark cycle. The welfare of all animals was continually assessed throughout all procedures.

LGI1 floxed mice (Chabrol et al., 2010) were bred with Cre lines to generate conditional KOs (cKO), such that Nav1.8 Cre line (LGI1^fl/Nav1.8(+/-)^) to target nociceptors and the HoxB8 Cre line (LGI1^fl/Hoxb^ ^(+/-)^) (Stirling et al., 2005; Witschi et al., 2010) to target all DRG neurons as well as spinal neurons. These mice were maintained on a C57BL/6J strain background, both male and female animals were included in the analysis and used aged between 6-16 weeks. Genotyping of offspring was performed by PCR of genomic DNA using LGI1 forward and reverse primers. Experiments are reported according to the ARRIVE guidelines.

### Cell culture

Adult male and female mice of 4-8 weeks of age were sacrificed in a CO2 chamber. The spinal column was rapidly removed and bisected to reveal the DRG. DRG were taken from all levels and placed directly into Hanks’ Balanced Salt solution (HBSS without Ca^2+^ and Mg^2+^, Invitrogen). DRG were then subjected to enzymatic digestion using collagenase II (12mg/ml, Worthington) and dispase II (14mg/ml, Roche) diluted in HBSS for 1.5 hours at 37°C. DRGs were then washed in HBSS and mechanically dissociated using fire polished pipettes. Dissociated cells were suspended in culture medium (Neurobasal medium, 2% B27, 1% GlutaMAX™, GIBCO, 1% antibiotic/antimycotic (ThermoFisher Scientific)) supplemented with mouse NGF (50ng/μl, Peprotech) and GDNF (10ng/ml, Peprotech), and plated on to 13 mm coverslips precoated with laminin (R&D Systems) and poly-D Lysine (BD biosciences) before being incubated at 37°C. After allowing the cells to attach to the coverslips for 2 h (at 37°C), the wells were flooded with the above-mentioned cell medium, and the primary culture was then left at 37°C overnight.

### Behavioral studies

At the start of each set of behavioral experiments, mice were acclimatised to the testing equipment and baseline values were obtained by averaging data from 3-4 sessions, and testing was performed at a consistent time of day. All tests were performed in the same room by the same experimenter, who was blind to all animal genotypes until after the behavioural analysis was complete and handled the mice in a random order. In total 21 LGI1^fl/Nav1.8(-/-)^ mice and 29 LGI1^fl/Nav1.8(+/-)^ littermates were used in behavioral studies (20 males and 30 females). In total, 25 LGI1^fl/Hoxb(-/-)^ mice and 31 LGI1^fl/Hoxb(+/-)^ littermates were used in behavioral studies (35 males and 21 females).

#### Von Frey

Mechanical sensitivity was assessed by placing mice in a Perspex box situated on top of a wire mesh. Calibrated Von Frey hairs (Ugo Basile) were then applied to the plantar surface of the hind paw and a reflex withdrawal response was used to calculate the 50% withdrawal threshold (Chaplan et al., 1994). Using a range of stimuli, we first applied the middle weight, 0.6 g, where a negative response increased the next weight applied, and a positive one – decreased it. The resulting pattern of responses was used to select a constant k and determine the final 50% withdrawal threshold. For spared nerve injury experiments mice were test for mechanical sensitivity over the course of the injury (days 3, 7, 14, 21, 28).

#### Brush

Dynamic mechanical allodynia was assessed using a recently published protocol (Cheng et al., 2017). A small paintbrush (5/0, The Art Shop) was modified by trimming the tip to make it blunt. This was used to gently stroke the plantar surface of the paw. A scoring system was used as follows to determine a dynamic allodynia score, (0) (a nonpainful response) lifting of the paw for less than 1 s, (1) sustained lifting of the paw or a single flinch, (2) lateral paw lift above the level of the body or a startle like jump and (3) multiple flinching responses or licking of the affected paw.

#### Pin Prick

The response to a clear noxious mechanical stimulus was also assessed using the pin prick test as previously described (Arcourt et al., 2017). A dissecting pin was attached to a 1 g Von Frey filament and applied to the plantar surface of the hind paw to elicit a rapid withdrawal reflex. The latency to withdraw was recorded using GoPro at 240 fps and analyzed using the video editing program Avidemux. The latency to withdraw from the pinprick analysed by an investigator blind to treatment groups. Three measurements were taken for each hind paw per trial and plotted latency represents the average of both paws.

#### Hargreaves

Thermal sensitivity was assessed using the Hargreaves method (Hargreaves et al., 1988). Here using the Hargreaves apparatus (Ugo Baslie) a radiant heat source was applied to the plantar surface of the hind paw and the latency to withdrawal was used to determine heat sensitivity threshold. Three measurements were taken for each hind paw and the averaged latency to withdraw was measured.

#### Hot Plate

Response to a suprathreshold heat stimulus was measured using the hot plate (Ugo Basile) assay. A metallic plate was set so that the surface temperature was at 53°C. Mice were chosen at random from their home cage and then placed onto the plate and the latency until a response, in this case shaking, licking, or biting of the paw, was measured.

#### Cold preference test

To assess cold sensitivity a thermal preference paradigm was used. The thermal preference equipment (Ugo Basile) consisted of two plates with a small connecting bridge. The plates were set at either 16°C or room temperature. Mice were chosen at random from their home cage and then assessed over a 10-minute period and the percentage of time spent at 16°C was calculated. For spared nerve injury experiments mice were test for thermal sensitivity over the course of the injury (days 6,13, 20, 27).

#### Formalin test

mice received an intraplantar injection of 20 μl of 5% formalin diluted from formaldehyde solution (5% v/v from 37% stock formaldehyde solution (Sigma)) in sterile saline. Mice were placed in a Perspex cylinder and the duration of pain-related behavior, biting/licking/paw lifting, was recorded over a 60-minute period while the experimenter left the room. The behavioral response is biphasic and therefore further comparisons were made by pooling data in the first (0-15mins) and second (15-60mins) phases.

### Spared Nerve Injury (SNI)

Mice were anesthetized using isoflurane inhalation and the left sciatic nerve and peripheral branches: common peroneal, tibial and sural nerves were exposed. The common peroneal and sural nerves were ligated, leaving the tibial nerve intact. The ligated nerves were transected distally, and a 2 mm section was removed to prevent nerve regeneration. The skin incision was then closed with 2-3 external stitches and appropriate post-operative care and analgesics given (local 2 mg/kg Marcain, AstraZeneca and systemic 5 mg/kg Rimadyl, Pfizer). Animals were used for behaviour or histology at least 5 weeks post-surgery. All behavioral measurements were taken in awake and unrestrained mice of both sexes. Von Frey, brush and cold preference experiments were performed as explained above. The experimenter was blind to all mouse genotypes.

### Conditioned Place Preference

To test for ongoing neuropathic pain, a three-day conditioning protocol using a biased chamber assignment was used for conditioned place preference (CPP) testing. The custom 3-chamber CPP apparatus consisted of two conditioning side chambers connected by a centre chamber (three chambered San Diego Instruments - dimensions of side chamber: 17cm W x 15cm D x 20cm H; dimensions of central chamber: 7cm W x 15cm D x 20cm H. Plexiglass inserts x2 for each central chamber – external dimensions: 6.5cm W x 2.5cm D x 17.5cm H). Isolation chamber (San Diego Instruments) – external dimensions: 60cm W x 29.5cm D x 50.5cm H). Mice were able to discriminate between chambers using visual (vertical black-and-white striped walls versus black-and-white spot walls) and sensory (strawberry versus vanilla scent) cues. On day 1 (acclimatisation, 4-5 weeks after tSNI surgery), mice had free access to explore all chambers for 30 minutes. On days 2 and 3 (preconditioning), mice were again allowed to freely explore for 30 min whilst their position was recorded using an infrared camera and AnyMaze 7.16 software (Stoelting, USA). To avoid pre-existing chamber bias, mice spending more than 80% or less than 5% of time in either side chamber during preconditioning were excluded. For conditioning (days 4 to 6), each morning, mice received i.p. vehicle injection (saline), were returned to their home cage for 5 min, then confined to their preferred side chamber for 30 min. Four hours later, mice received i.p. gabapentin (100 mg/kg), were returned to their home cage for 5 min, and then placed in their non-preferred chamber for 30 min. On test day (day 7), mice could freely explore all chambers whilst their position was recorded, as during pre-conditioning, for 30 min. Difference scores were calculated as the time spent in each chamber on test day minus the mean time spent during pre-conditioning.

### Histology

Tissue preparation: For immunohistochemistry and in situ hybridization studies, mice were overdosed with pentobarbital and transcardially perfused, initially with sterile saline and then 20mls of 4% paraformaldehyde (PFA, 0.1M Phosphate buffer (PB)). Once dissected the DRG were post-fixed in 4% PFA for 1.5 hours at RT and spinal cord overnight at 4°C. All tissue was dehydrated for cyroprotection in 30% sucrose (0.1M PB) at 4°C for at least 24 hours. Tissue was then embedded in optimal cutting temperature (OCT) medium (Tissue-Tek) and stored at −80. Tissue was then placed into a solution containing only 30% sucrose before being embedded in OCT. Tissue was sectioned onto Superfrost plus slides (VWR) using a cryostat. DRG sections were cut at 10 μm and spinal cord at 20 μm. The slides were then stored at −80.

### Immunohistochemistry (IHC)

DRG and spinal cord tissue sections were washed once in PBS and PBS Triton-X (0.3%), before being incubated overnight at room temperature with the respective primary antibodies diluted in PBS/Triton. Primary Ab was washed off in PBS triton-X and tissue was then incubated with secondary antibodies at RT for 2-4 hours (Alexa Fluor, Thermo Fisher Scientific). The tissue was then washed again and cover-slipped. Immunostaining was visualized using a confocal microscope (Axio LSM 700, Zeiss) and images acquired using the Zen black software.

### In situ hybridization (ISH)

Once cut, DRG and spinal cord sections were air-dried overnight and then stored in the −80°C. ISH was carried out using the RNAScope 2.5 RED chromogenic assay kit and by following the manufacturer’s instructions (Advanced Cell Diagnostics). Briefly, tissue sections were removed from the −80, allowed to equilibrate to RT and re-hydrated in PBS. Pre-treatment required a hydrogen peroxide step at RT; followed by a protease treatment in a hybridization oven at 40°C. Slides were then incubated with the target or control probes at 40°C for 2 hours. Slides were then incubated with an mRNA probe against LGI1 for 2 h at 40°C. Following probe incubation, slides were subjected to 6 rounds of amplification and the probe signal was developed via a reaction with fast red. To combine with IHC, tissue sections were then washed with PBS-Tx (0.3%) and subjected to the standard IHC protocol.

### Image Analysis

Analysis of the signal intensity for in situ hybridization studies on DRG was calculated using ImageJ software. In a single image of a section of either L4 or 5 DRG, neurons with a nucleus were circled and the percentage coverage of red signal for that cell profile area measurement was calculated. By eye the cell was then subpopulation defined by using primary antibodies against NF200, IB4, CGRP or TH and the appropriate secondary antibody. For each marker at least 3 sections were imaged per animal. On each image a background reading was taken from an area of tissue not containing neuronal cell bodies. An average of this was calculated for each animal and any cell that had percentage coverage of greater than 2 standard deviations from the background mean were defined as being positive for LGI1 mRNA. For spinal cord sections, LGI1 mRNA positive cells were defined as those containing 4 or more red dots. For analysis of neuronal activity following formalin (5%) injection, LGI1^fl/Hoxb^ ^(+/-)^ and control mice were perfused two hours post injection and tissue was incubated with a primary antibody against c-fos. NeuN was used to mark neuronal cell bodies. The average number of c-fos positive neurons in the dorsal horn was calculated from 3-5 sections per animal for both ipsilateral and contralateral sides. For the quantification of DRG neuron subpopulations in mice, 3 sections of the L4 DRG were imaged and used for each animal with > 100 cells counted for each subpopulation. The number of cells for each marker is shown as a percentage of the sum of NF200, IB4 and CGRP positive cells. For analysis of PAX2 positive inhibitory interneurons in mice, tissue was incubated in primary antibody against PAX2 and 3 sections from the L4 segment of the mouse spinal cord were scanned. All quantification was performed with the experimenter blind to treatment groups.

### Confocal imaging

Images were captured on the Zeiss LSM 700, using 405nm, 488nm and 546nm diode lasers. One z stack per section was taken at an interval of 0.1 μm across the central portion of the superficial dorsal horn, covering lamina 1-3 dorsoventrally. Approximately 50 optical sections per image were taken for analysis. All image processing and analysis was performed in ImageJ.

### RNA isolation and cDNA synthesis

Mice were culled using a CO2 chamber. Dissected DRGs were immediately frozen on liquid nitrogen and stored at −80. RNA was isolated using a combination of TriPure (Roche) and a High Pure RNA tissue kit (Roche). Briefly, tissue was homogenized in Tripure using a handheld homogenizer (Cole-Parmer) treated with chloroform and then subjected to column purification before being eluted in RNase free water. Synthesis of cDNA was carried out using Transcriptor reverse transcriptase (Roche), random hexamers (Invitrogen) and dNTPs (Roche).

### Quantitative Real Time PCR

For analysis of mRNA expression using SYBR green cDNA (5ng) and primers (0.5μM) were mixed with LightCycler 480 SYBR Green Master (Roche) in a 1:1 ratio and added to white 384 well plates (Roche). Plates were run on a 45-cycle protocol using the LC 480 II system (Roche).

Primers were designed using Primer-BLAST (https://www.ncbi.nlm.nih.gov/tools/primer-blast/). Primer efficiency and specificity were validated before experimental use. Gene expression for each target primer was normalized against 3 reference genes (18 s, GAPDH and HPRT1) using the delta delta CT method.

### Electrophysiology

Patch clamping of DRG neurons in vitro: Whole-cell patch clamp recordings were performed at room temperature (22°) using an Axopatch 200B amplifier and Digidata 1550 acquisition system (Molecular Devices). Data were low-pass filtered at 2 kHz and sampled at 10 kHz. Series resistance was compensated 70%–90% to reduce voltage errors. Patch pipettes (2–4MU) were pulled from filamental borosilicate glass capillaries (1.5 mm OD, 0.84 mm ID; World Precision Instruments) Patch pipettes were filled with internal solution containing (mM): 130 KCl, 1 MgCl2, 5 MgATP, 10 HEPES, and 0.5 EGTA; pH was adjusted to 7.3 with KOH and os- molarity set to 305 mOsm. Extracellular solution contained (mM): 140 NaCl, 4.7 KCl, 1.2 MgCl2, 2.5 CaCl2, 10 HEPES and 10 glucose; pH was adjusted to 7.3 with NaOH and osmolarity was set to 315 mOsm. Resting membrane potential was assessed in bridge mode, while firing properties were assessed in current clamp mode. Input resistance (RInput) was calculated from the voltage deflections caused by increasing (D20 pA) hyperpo- larising current pulses. To determine rheobase, cells were depolarised from a holding potential of -60 mV by current steps (50 ms) of increasing magnitude (D25 pA) until an action potential was generated. Repetitive firing was assessed by 500 ms depolar- ising current steps of increasing magnitude (50pA). Data were analyzed by Clampfit 10 software (Molecular Devices)

### Calcium Imaging of DRG

Coverslips were incubated for 45-90 min at 37°C with 1 μM Fura-2AM (Invitrogen) in Neurobasal medium supplemented as above. After incubation, coverslips were transferred to artificial extracellular fluid (140 mM NaCl, 5 mM KCl, 2 mM CaCl2, 1 mM MgCl2, 10 mM D-Glucose, 10 mM HEPES in distilled water). Coverslips were imaged every 1s with 4x4 binning for 340 s on a Zeiss inverted fluorescence microscope with a 10x objective, dichroic LP 409 mirror, BP 340/30 and BP 387/15 excitation and 510/90 emission filters. ZEN Blue software was used for image acquisition and selection of regions of interest (ROIs).

ECF was perfused continuously over the cells. After 300s of baseline recording, the cells were perfused with 10μM ATP for 30 s. This was followed by a 180s washout and then 1μM capsaicin treatment was added for 30s followed by 180s washout. Neurons were then identified by their responsiveness to 50 mM KCL (30s) and followed by a 30s washout.

Data were then analysed using a Matlab script. The amplitude and rate threshold values (at 0.15 and 0.01 for fura2 imaging, respectively) were detected using Excel and a Matlab script. If the maximum amplitude of the calcium trace relative to the baseline (average from 0s to start of first stimulus) is greater than the chosen threshold (0.15units) and the rate of change crosses the chosen threshold (here 0.01) at least once during the analysis period, a response will be recorded. Only neurons that had a KCL response were considered for analysis. The percentage of cells responding to a certain stimulus was then calculated.

### Quantification and Statistical Analysis

Data is shown as the mean ± SEM, unless otherwise stated. A Student’s t test was used to compare the mean of two groups and when data was not normally distributed a non-parametric test was applied (Mann-Whitney). A one-way ANOVA was used when more than two groups existed. For behavioral studies over time, a repeated-measures two-way ANOVA was used with posthoc Tukey analysis. For calcium imaging a one-way ANOVA was performed. For patch clamp experiments multiple Mann-Whitney tests were used for repetitive firing assays on DRG cells and a two-way ANOVA with posthoc Tukey analysis for assessing excitability parameters. Sample sizes for each experimental can be found in the figure legends. Significance for all experiments was placed at p < 0.05. Statistical tests were carried out with Sigmaplot.

## Key Resources Table

| Reagent or Resource | Source | Identifier |
| --- | --- | --- |
| **Antibodies** |  |  |
| NeuN (1:500, Chicken) | Merck Millipore | Abn91 |
| IB4(1:50,Streptavidin conjugated) | Sigma-Aldrich | L2140 |
| CGRP (1:250, Sheep) | Enzo | Ca1137 |
| NF200 (1:1000, Rabbit) | Merck Millipore | ABN76 |
| Pacific Blue (1:100) | Thermo Fischer Scientific | Alexa Fluor (S11222) |
| Alexa Fluor 488 (1:500) | Thermo Fischer Scientific | Alexa Fluor (A110008) |
| Alexa Fluor 546 (1:500) | Thermo Fischer Scientific | Alexa Fluor (A11040) |
| Neurotrace (1:500) | Thermo Fischer Scientific | N21482 |
| PAX2 (1:500, rabbit) | Thermo Fischer Scientific | 71-6000 |
| C-FOS (1:500, rabbit) | Synaptic Systems | 226 008 |

## Acknowledgements

The authors thank John Wood for providing the Nav1.8Cre mouse line, Hanns Ulrich Zeilhofer for providing the HoxB Cre line, and Stephanie Baulac for providing the LGI1 floxed line.

## Data and materials availability

All data and materials will be made available upon reasonable request to the corresponding author(s).

