## Supplementary figures and tables for "A role for LGI1 in regulating pain sensitivity"

### Figure S1

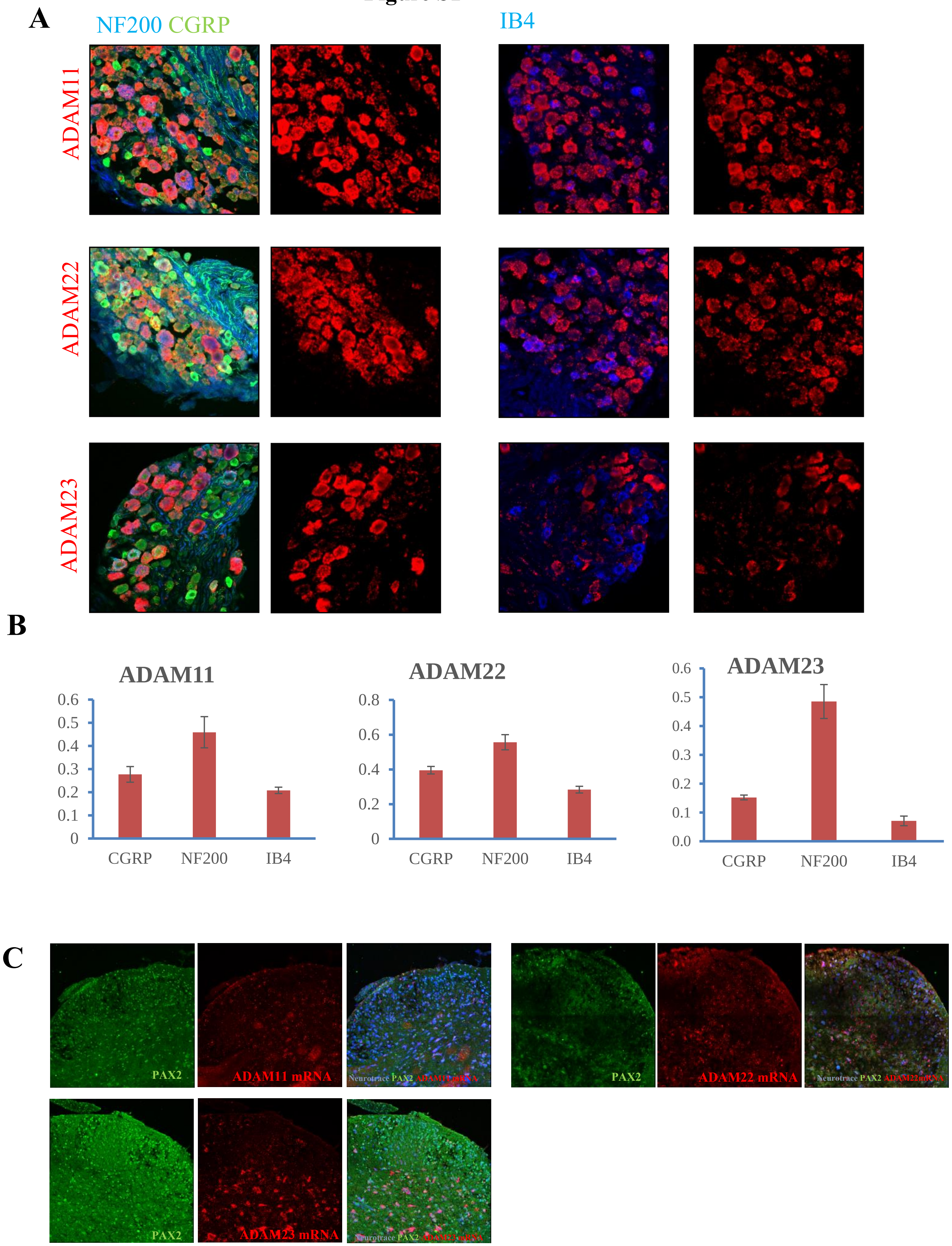
A
NF200 CGRP
IB4
ADAM11
ADAM22
ADAM23
B
ADAM11
0.6
0.5
0.4
0.3
0.2
0.1
0
CGRP
NF200
IB4
ADAM22
0.8
0.6
0.4
0.2
0
CGRP
NF200
IB4
ADAM23
0.6
0.5
0.4
0.3
0.2
0.1
0.0
CGRP
NF200
IB4
C
PAX2
ADAM11 mRNA
Neurotrace PAX2 ADAM11 mRNA
PAX2
ADAM22 mRNA
Neurotrace PAX2 ADAM22 mRNA
PAX2
ADAM23 mRNA
Neurotrace PAX2 ADAM23 mRNA

**Figure S2**

**A** ***DRG***

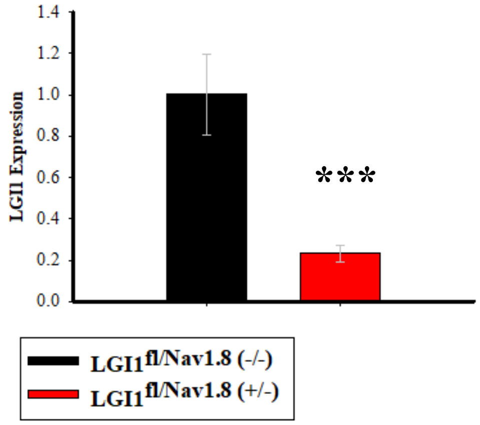


**B**

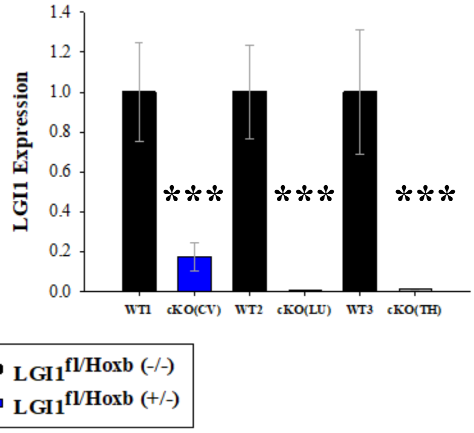


***Spinal cord & Brain***

**C**

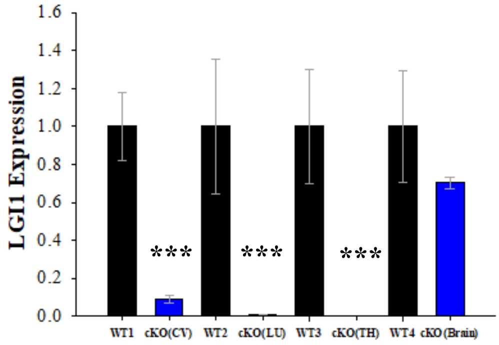

**Figure S3**

**A**

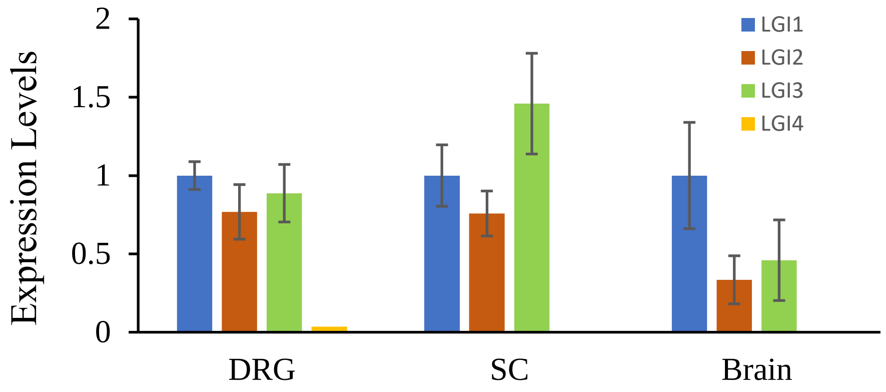


***DRG:***

**B**

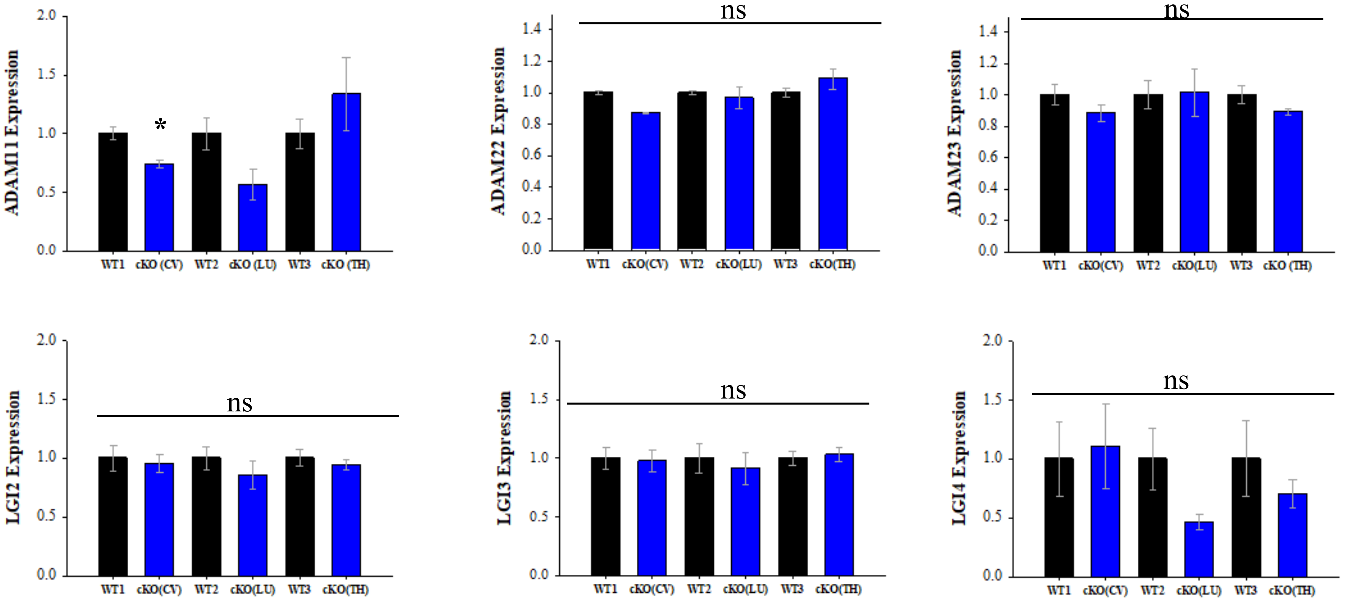


**C** ***Spinal Cord & Brain:***

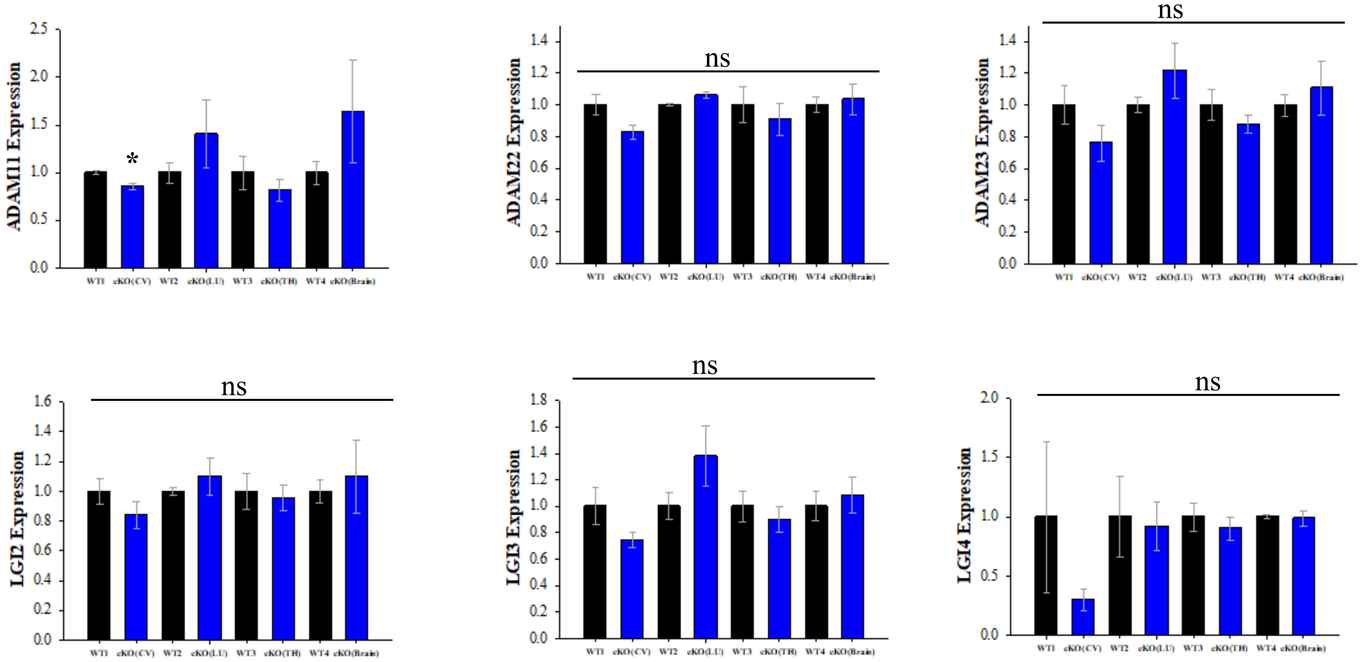

### Figure S4

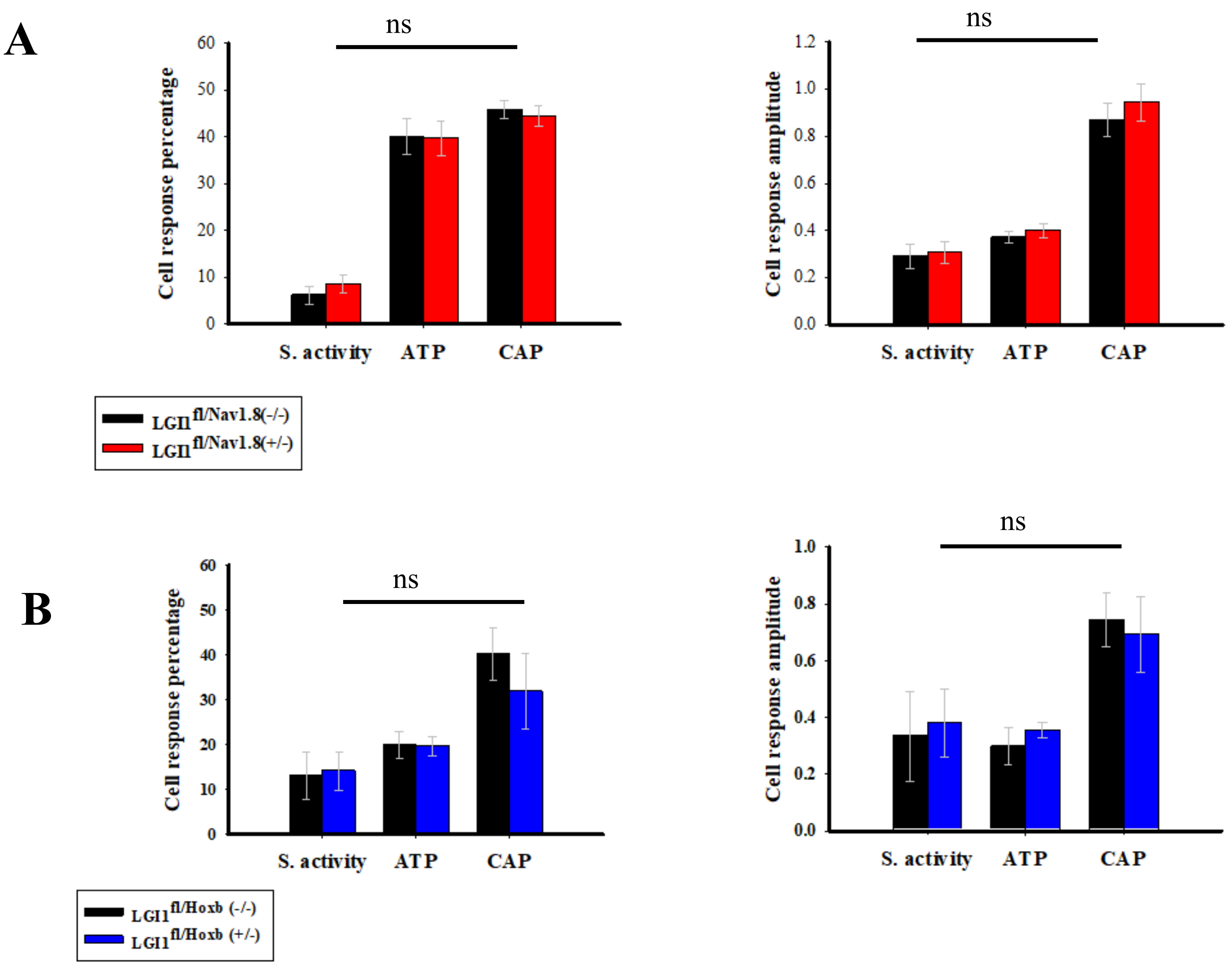

A
ns
Cell response percentage
0
10
20
30
40
50
60
S. activity
ATP
CAP
LGI1fl/Nav1.8(-/-)
LGI1fl/Nav1.8(+/-)
ns
Cell response amplitude
0.0
0.2
0.4
0.6
0.8
1.0
1.2
S. activity
ATP
CAP
B
ns
Cell response percentage
0
10
20
30
40
50
60
S. activity
ATP
CAP
LGI1fl/Hoxb (-/-)
LGI1fl/Hoxb (+/-)
ns
Cell response amplitude
0.0
0.2
0.4
0.6
0.8
1.0
S. activity
ATP
CAP

### Figure S5

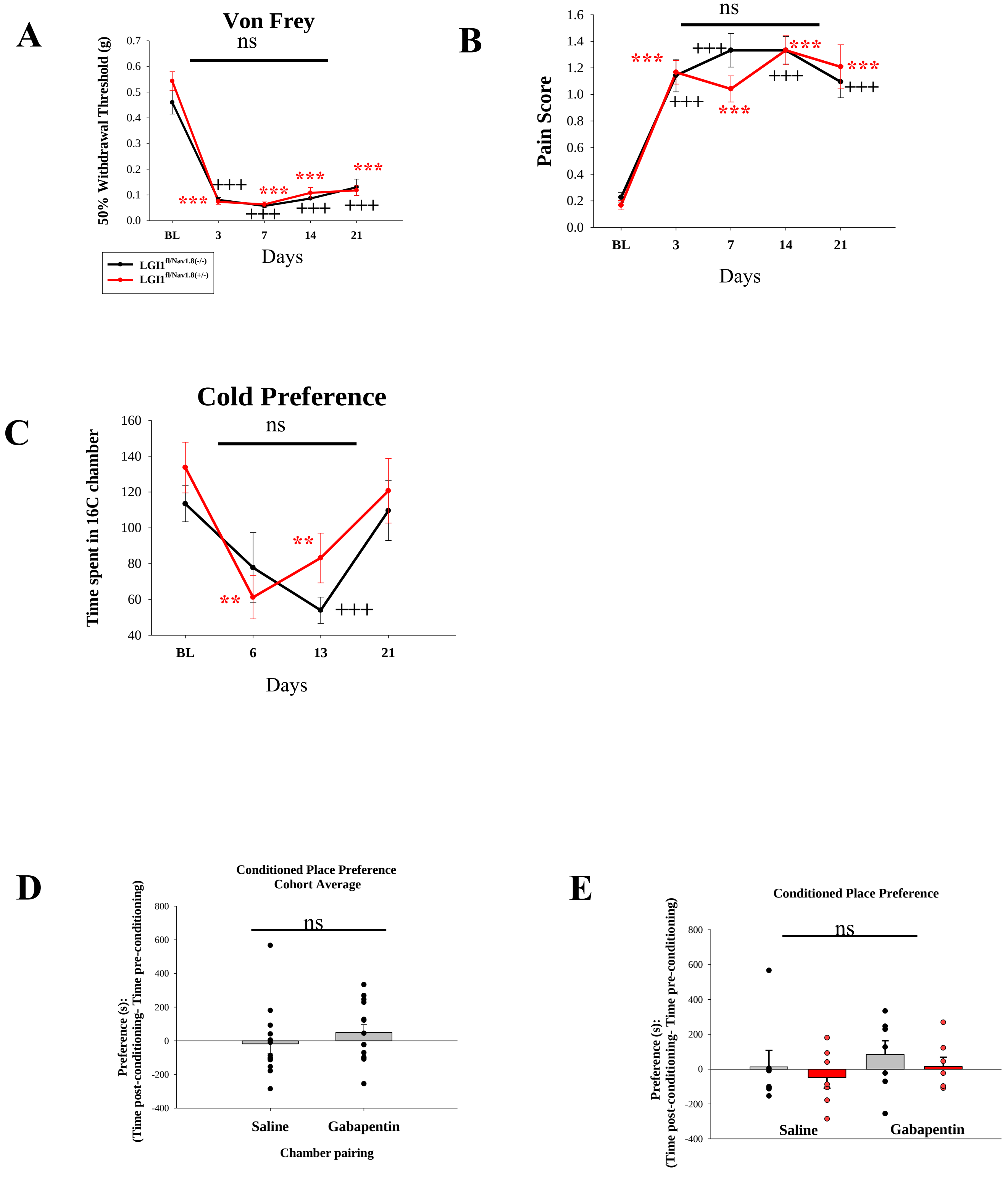

A
Von Frey
ns
50% Withdrawal Threshold (g)
0.0
0.1
0.2
0.3
0.4
0.5
0.6
0.7
BL
3
7
14
21
Days
***
+++
LGI1 fl/Nav1.8(-/-)
LGI1 fl/Nav1.8(+/-)
B
Brush
ns
Pain Score
0.0
0.2
0.4
0.6
0.8
1.0
1.2
1.4
1.6
BL
3
7
14
21
Days
***
+++
C
Cold Preference
ns
Time spent in 16C chamber
40
60
80
100
120
140
160
BL
6
13
21
Days
**
+++
D
Conditioned Place Preference
Cohort Average
ns
Preference (s):
(Time post-conditioning- Time pre-conditioning)
800
600
400
200
0
-200
-400
Saline
Gabapentin
Chamber pairing
E
Conditioned Place Preference
ns
Preference (s):
(Time post-conditioning- Time pre-conditioning)
800
600
400
200
0
-200
-400
Saline
Gabapentin

**Fig.S1: Expression levels of ADAMs proteins in DRG and spinal cord. (A)** ADAMs 11, 22 and 23 are known to act as LGI1 binding partners in the CNS and these interactors are expressed by DRG neurons. Representative images of ISH for ADAMs mRNA in DRG sections stained for molecular markers of DRG neuron subtypes: IB4, CRGP, and NF200. **(B)** Quantification of signal intensity for each probe. Data shown as mean±SEM (n=3-4 mice). **(C)** ADAM11,22,23 expression in the mouse spinal cord. Representative images of ISH for ADAMs in spinal cord showing also neurotrace-positive neurons co-localizing with ADAMs & Pax2 in the superficial laminae and in the deeper laminae of the spinal cord.

**Fig.S2: (A)** qPCR data showing that the expression level of LGI1 decreased by 70% in the LGI1 knockout mice from the Nav 1.8-Cre mouse line (n=3, *** p<0.001, t test). **(B-C)**, qPCR data showing that LGI1 expression was significantly reduced at all DRG and spinal levels (cervical (CV), lumbar (LU), thoracic (TH) in HoxB Cre mouse line (n=3, ***p<0.001, One WAY ANOVA)

**Fig. S3: (A)** qPCR data showing the expression levels of LGI1,2,3,4. LGI2 and 3 had a similar expression pattern and profile to that of LGI1 in both the DRG and spinal cord, and lesser expression levels were detected in the brain in comparison to LGI1 expression levels. Slight expression levels were detected for the LGI4 protein in all three types of tissues relative to LGI1 expression levels (n=3, mean±SEM). **(B-C)** qPCR data showing expression levels of the ADAM 11,22,23 and the proteins LGI2, 3, and 4 in the mouse HoxB Cre line. qPCR data indicate that the expression levels of the ADAMs proteins were mostly similar between the cKO and WT mice in the DRG, spinal cord, and brain. ADAM11 protein displayed a slight significant decrease in expression levels on the cervical level of the DRG and spinal cord of the cKO mice compared to the WT (cervical (CV), lumbar (LU), thoracic (TH), *p<0.05, n=3, One way ANOVA, mean±SEM). The expression levels of LGI2, 3 and 4 were all similar between the two genotypes in the DRG, spinal cord, and brain.

**Fig.S4: Calcium imaging results assessing spontaneous activity in DRG neurons from Nav 1.8 Cre line and HoxB Cre lines.** Cell response and amplitude were also assessed after 1μM capsaicin (CAP) or 10μM ATP treatment. Neurons were identified by their responsiveness to 50 mM KCl. **(A)** No significant difference in cell response percentage and amplitude was observed between genotypes for the Nav 1.8 Cre line for the three tested conditions (n=3, One way ANOVA, mean±SEM). **(B)** Similar results were also obtained for the HoxB Cre line for the percentage of cells responding to ATP or capsaicin treatments as well for any spontaneous activity (n=6, One way ANOVA, mean±SEM).

**Fig.S5: The effects of the genetic removal of LGI1 in the context of spared nerve injury (tibial) model (tSNI) were assessed in Nav 1.8 Cre line. (A-C),** All mice received surgery and mechanical and thermal sensitivity assessed over 3-4 weeks. $LGI1^{fl/Nav1.8(-/-)}$ (n=7) and $LGI1^{fl/Nav1.8(+/-)}$ (n=8) mice became hypersensitive to mechanical stimuli and cold stimuli by day 3 and 6, respectively, but no significant differences between genotypes were found (* vs baseline (BL), + vs BL, **p<0.01, *** and +++ p<0.001, two-way repeated measures ANOVA posthoc Tukey, mean±SEM). **(D-E)** ongoing neuropathic pain was investigated in mice following tSNI using the conditioned place preference test with no significant differences in the CPP difference scores between genotypes were observed ($LGI1^{fl/Nav1.8(-/-)}$ (n=7) and $LGI1^{fl/Nav1.8(+/-)}$ (n=8), p>0.05, One Way ANOVA, mean±SEM).

**Table S1**: Quantification of signal intensity for LGI1 and ADAMs 11,22, 23 probes in DRG neuron subtypes. Related to Fig.1 and Fig.S1.

| Gene Name | CGRP | NF200 | IB4 |
|---|---|---|---|
| LGI1 | 0.32±0.09 | 0.32±0.09 | 0.15±0.07 |
| ADAM11 | 0.28±0.03 | 0.46±0.07 | 0.21±0.01 |
| ADAM22 | 0.39±0.02 | 0.56±0.04 | 0.28±0.02 |
| ADAM23 | 0.15±0.06 | 0.485±0.02 | 0.09±0.03 |

**Tabe S2:** qPCR analysis of gene expression of LGI1 and ADAMs 11, 22, 23 in DRG, spinal cord and brain. Related to Fig.1C.

| Gene Name | DRG | Spinal Cord | Brain |
|---|---|---|---|
| LGI1 | 1±0.07 | 1.09±0.02 | 1.31±0.07 |
| ADAM11 | 1±0.15 | 0.05±0.01 | 0.09±0.04 |
| ADAM22 | 1±0.09 | 0.41±0.08 | 0.43±0.04 |
| ADAM23 | 1±0.07 | 0.64±0.03 | 0.47±0.02 |

**Table S3:** qPCR analysis of gene expression of LGI1, 2, 3,4 in DRG, spinal cord and brain. Related to Fig.S3A.

| Gene Name | DRG | Spinal Cord | Brain |
|---|---|---|---|
| LGI1 | 1±0.09 | 1±0.19 | 1±0.34 |
| LGI2 | 0.77±0.17 | 0.76±0.14 | 0.33±0.15 |
| LGI3 | 0.89±0.18 | 1.46±0.32 | 0.46±0.26 |
| LGI4 | 0.04±0.01 | 0.003±0.001 | 0.002±0.0008 |

**Table S4:** qPCR analysis of gene expression of ADAMs 11, 22, 23 and LGI2,3,4 in DRG, spinal cord and brain in HoxB Cre line mice. Related to Fig.S3B,C.

| Gene name | DRG cervical | | DRG lumbar | | DRG thoracic | |
|---|---|---|---|---|---|---|
| | LGI1fl/HoxB(-/-) | LGI1fl/HoxB(+/-) | LGI1fl/HoxB(-/-) | LGI1fl/HoxB(+/-) | LGI1fl/HoxB(-/-) | LGI1fl/HoxB(+/-) |
| ADAM11 | 1±0.05 | 0.74±0.04 | 1±0.14 | 0.57±0.13 | 1±0.12 | 1.33±0.31 |
| ADAM22 | 1±0.01 | 0.87±0.01 | 1±0.0.01 | 0.97±0.07 | 1±0.03 | 1.09±0.07 |
| ADAM23 | 1±0.06 | 0.89±0.05 | 1±0.09 | 1.01±0.15 | 1±0.05 | 0.89±0.02 |
| LGI2 | 1±0.11 | 0.95±0.07 | 1±0.09 | 0.85±0.12 | 1±0.07 | 0.94±0.05 |
| LGI3 | 1±0.09 | 0.98±0.09 | 1±0.13 | 0.92±0.14 | 1±0.06 | 1.03±0.06 |
| LGI4 | 1±0.32 | 1.10±0.36 | 1±0.26 | 0.47±0.06 | 1±0.32 | 0.70±0.12 |

| Gene name | Spinal cord cervical | | Spinal cord lumbar | | Spinal cord thoracic | |
|---|---|---|---|---|---|---|
| | LGI1fl/HoxB(-/-) | LGI1fl/HoxB(+/-) | LGI1fl/HoxB(-/-) | LGI1fl/HoxB(+/-) | LGI1fl/HoxB(-/-) | LGI1fl/HoxB(+/-) |
| ADAM11 | 1±0.02 | 0.85±0.03 | 1±0.11 | 1.40±0.36 | 1±0.18 | 0.81±0.12 |
| ADAM22 | 1±0.07 | 0.83±0.05 | 1±0.01 | 1.06±0.02 | 1±0.11 | 0.91±0.10 |
| ADAM23 | 1±0.12 | 0.76±0.11 | 1±0.05 | 1.22±0.17 | 1±0.09 | 0.88±0.06 |
| LGI2 | 1±0.08 | 0.84±0.09 | 1±0.03 | 1.10±0.12 | 1±0.12 | 0.96±0.09 |
| LGI3 | 1±0.14 | 0.74±0.06 | 1±0.10 | 1.38±0.23 | 1±0.12 | 0.90±0.09 |
| LGI4 | 1±0.64 | 0.29±0.09 | 1±0.34 | 0.92±0.21 | 1±0.12 | 0.89±0.09 |

| Gene Name | Brain | |
|---|---|---|
| | LGI1fl/HoxB(-/-) | LGI1fl/HoxB(+/-) |
| ADAM11 | 1±0.12 | 1.64±0.54 |
| ADAM22 | 1±0.05 | 1.04±0.09 |
| ADAM23 | 1±0.07 | 1.11±0.17 |
| LGI2 | 1±0.08 | 1.10±0.25 |
| LGI3 | 1±0.11 | 1.08±0.14 |
| LGI4 | 1±0.02 | 0.99±0.07 |

**Table S5:** Behavior analysis for Nav1.8 Cre line mice. Related to Fig. 3A

| Behaviour test | LGI1fl/Nav1.8(-/-) | No. of mice | LGI1fl/Nav1.8(+/-) | No.mice |
|---|---|---|---|---|
| Pin Prick | 121.12±13.40 | 6 | 118.32±13.17 | 12 |
| Hot Plate 53 | 7.96±0.63 | 8 | 7.53±0.52 | 9 |
| Von Frey | 0.50±0.04 | 14 | 0.55±0.04 | 21 |
| Hargreaves | 10.2±0.69 | 14 | 8.4±0.42 | 21 |

**Table S6:** Behavior analysis for HoxB Cre line mice. Related to Fig. 3B

| Behaviour test | LGI1fl/HoxB(-/-) | No. of mice | LGI1fl/HoxB(+/-) | No.mice |
|---|---|---|---|---|
| Pin Prick | 270.17±50.47 | 19 | 287.79±40.47 | 22 |
| Hot Plate 53 | 8.51±0.47 | 19 | 8.74±0.38 | 22 |
| Von Frey | 0.61±0.02 | 25 | 0.53±0.02 | 31 |
| Hargreaves | 9.81±0.39 | 19 | 9.65±0.33 | 22 |

**Table S7:** Calcium imaging results looking at cell response and amplitude from Nav 1.8 Cre and HoxB Cre mouse lines in response to capsaicin and ATP, and assessing spontaneous activity. Related to Fig. S4

| | Cell response percentage | | Cell response amplitude | |
|---|---|---|---|---|
| | LGI1fl/Nav1.8(-/-) | LGI1fl/Nav1.8(+/-) | LGI1fl/Nav1.8(-/-) | LGI1fl/Nav1.8(+/-) |
| Spontaneous activity | 6.17±1.88 | 8.63±1.99 | 0.29±0.05 | 0.31±0.05 |
| ATP | 40.08±3.82 | 39.79±3.74 | 0.37±0.02 | 0.39±0.03 |
| Capsaicin | 45.83±1.95 | 44.40±2.17 | 0.87±0.07 | 0.94±0.08 |

| | Cell response percentage | | Cell response amplitude | |
|---|---|---|---|---|
| | LGI1fl/HoxB(-/-) | LGI1fl/HoxB(+/-) | LGI1fl/HoxB(-/-) | LGI1fl/HoxB(+/-) |
| Spontaneous activity | 13.14±5.28 | 14.18±4.32 | 0.33±0.16 | 0.38±0.12 |
| ATP | 19.98±2.99 | 19.69±2.13 | 0.29±0.07 | 0.36±0.03 |
| Capsaicin | 40.22±5.93 | 31.88±8.43 | 0.74±0.09 | 0.69±0.14 |

**Table S8:** Biophysical properties in cultured DRG neurons from Nav 1.8 Cre and HoxB Cre mouse lines. Related to Fig. 3.

| | Small (<25µm) | | Medium (25-35 µm) | |
|---|---|---|---|---|
| | **LGI1fl/Nav1.8(-/-)** | **LGI1fl/Nav1.8(+/-)** | **LGI1fl/Nav1.8(-/-)** | **LGI1fl/Nav1.8(+/-)** |
| R Input (MΩ) | 284.81±16.25 | 301.07±25.15 | 109.49±8.38 | 96.88±19.93 |
| RMP (mV) | -49.86±0.74 | -49.86±0.91 | -56.45±1.25 | -57.76±1.10 |
| Rheobase (pA) | 272.21±21.14 | 261.07±30.25 | 779.16±90.36 | 1126.85±177.33 |
| No. of cells | 39 | 35 | 17 | 20 |
| | Small (<25µm) | | Medium (25-35 µm) | |
| | **LGI1fl/HoxB(-/-)** | **LGI1fl/HoxB(+/-)** | **LGI1fl/HoxB(-/-)** | **LGI1fl/HoxB(+/-)** |
| R Input (MΩ) | 270.88±15.80 | 285.29±19.33 | 86.69±9.71 | 65.22±5.86 |
| RMP (mV) | -48.86±0.69 | -49.17±0.66 | -55.65±1.23 | -57.24±0.85 |
| Rheobase (pA) | 293.41±24.69 | 284.50±20.27 | 1259.56±209 | 1499.28±180.16 |
| No. of cells | 47 | 46 | 23 | 23 |
